# EvoBind: *in silico* directed evolution of peptide binders with AlphaFold

**DOI:** 10.1101/2022.07.23.501214

**Authors:** Patrick Bryant, Arne Elofsson

## Abstract

Currently, there is no accurate method to computationally design peptide binders towards a specific protein interface using only a target structure. Experimental methods such as phage display can produce strong binders, but it is impossible to know where these bind without solving the structures. Using AlphaFold2 (AF) and other AI methods to distinguish true binders has proven highly successful but relies on the availability of binding scaffolds. Here, we develop EvoBind, an *in silico* directed-evolution platform based on AF that designs peptide binders towards an interface using only sequence information. We show that AF can distinguish between native and mutated peptide binders using the plDDT score and find that AF adapts the receptor interface structure to the binders during optimisation. We analyse previously designed minibinder proteins and show that AF can distinguish designed binders from non-binders. We compare ELISA ratios of different peptide binders and find the affinity can not be distinguished among binders, possibly due to varying binding sites and low AF confidence. We test the recovery of binding motifs and find that up to 75% of motifs are recovered. In principle, EvoBind can be used to design binders towards any interface conditioned on if AF can predict these. We expect that EvoBind will aid experimentalists substantially, providing a starting point for further laboratory analysis and optimisation. We hope that the use of AI-based methods will come to make binder design significantly cheaper and more accurate in tackling unmet clinical needs. EvoBind is freely available at: https://colab.research.google.com/github/patrickbryant1/EvoBind/blob/master/EvoBind.ipynb

## Introduction

To be able to design peptide binders towards specified protein interfaces is a highly coveted goal which would have a major impact on pharmaceutical development [1]. Even though peptide drugs constitute only 5% of the pharmaceutical market, there has been an 8% average annual increase of approved peptide drugs during the past six decades and global sales exceed $50 billion yearly [2]. Previously, the state-of-the-art for computer-aided binder design[3] relied on docking scoring programs which are now significantly outperformed by AlphaFold [4–6]. The resulting outcome was that only one in 100’000 designed sequences bind to their target structures [3]. Alternatively, experimental techniques such as directed evolution can be used together with classical machine learning [7,8] to improve the outcome coupled with phage display[9]. None of the experimental methods can be used to design a binder towards a certain interface region however, potentially resulting in binding optimisation towards unintended protein regions.

Recent developments in deep learning, mainly based on AlphaFold2 (AF) have shown great promise for structure prediction of protein-protein[4,5] and protein-peptide[10] interactions as well as protein design[11,12]. A recent protein design method, ProteinMPNN[13], improved further on these achievements and created proteins with significantly higher solubility compared to using AF alone. ProteinMPNN can also generate binder sequences with higher success rate than previously, although it relies on specifying the backbone trace of a binder structure[14]. Reevaluating previous designs [3] using AF as a scoring function reports experimental success rates of close to 90% in some cases compared to only 0-5% using physics based calculations [14]. These methods have been applied to design binders both with and without scaffolds and known binding motifs[15]. Other methods such as MaSIF utilise learned interaction potentials and protein scaffold search to design binders[16]. However, no method relying only on sequence information targeted towards a certain interface region exists, limiting the possibilities of binders to previously available structures.

By applying language models trained on all available protein sequences[17] it is possible to predict the direction of evolution [18]. The probabilities associated with these language model predictions have been shown to inform successful affinity maturation of antibodies from screening only 20 variants [19]. The recent success of computational methods for protein design and directed evolution sprouted the idea to build a framework for designing peptide binders towards specific protein interfaces that is purely computational. If a peptide of only 10 amino acids is to be designed, the number of possibilities are 20^10^ ≈ 1.02·10^13^ considering the standard amino acids. This vast number of possibilities is not searchable in a reasonable amount of time, and it is not known if AF can distinguish between truly binding and nonbinding sequences.

We explore the possibility of peptide binder sequence design based on AF using only the sequence of the target protein and specified interface residues. We create binders towards a target interface by updating the sequence based on several loss terms that capture the interaction potential and the overall ability of AF to predict the structure. Together, our approach results in an efficient guide to design peptide binders, with the possibility to design binders for user-specified regions of different proteins.

## Results and Discussion

Here, we outline the investigations performed. First, we analyse the impact of the number of recycles on the ability of AlphaFold2 (AF) to predict the structure of protein-peptide interactions and select an optimal setting for the following analyses. We show that we can distinguish between correct predictions with very high accuracy (ROC AUC=0.94) and continue to analyse the ability of AF to distinguish between mutated peptide sequences. We create a search strategy for the design of peptide binders and demonstrate the success of our approach. Subsequently, we analyse the structural adaptation of the receptor to the designed peptides, the possibility to find alternative binding sites, distinguish designed miniprotein binders, peptide binding affinity and recover binding motifs.

### Peptide structure prediction

AlphaFold2 (AF) is able to predict the structure of protein-peptide complexes using a multiple sequence alignment (MSA) to represent the target structure, and the single sequence for the peptide representation [10]. The success of this approach was however reported using an increased number of recycles (nine) and as the best of ten different models. If one aims to search for a sequence using AF as a target function, having many recycles is not effective and a top-10 approach is not realistic. Therefore, we evaluate the performance of AF on a set of 96 non-redundant peptides[10] using 1-10 recycles on top-1 predictions. Figure 1a shows the cumulative fraction of peptides below a certain threshold in the average interface (IF) RMSD for different numbers of recycles. The interface is defined as beta carbons (Cβs) within 8 Å between the peptide and its receptor protein and the IF RMSD was calculated after aligning the alpha carbons (CAs) between the predicted and native structures. We find that at above 8 recycles no improvement is observed and 12.5% of the models (n=12 out of 96) are within 2 Å RMSD. At one recycle, only six models are correct at 2 Å.

**Figure 1.**
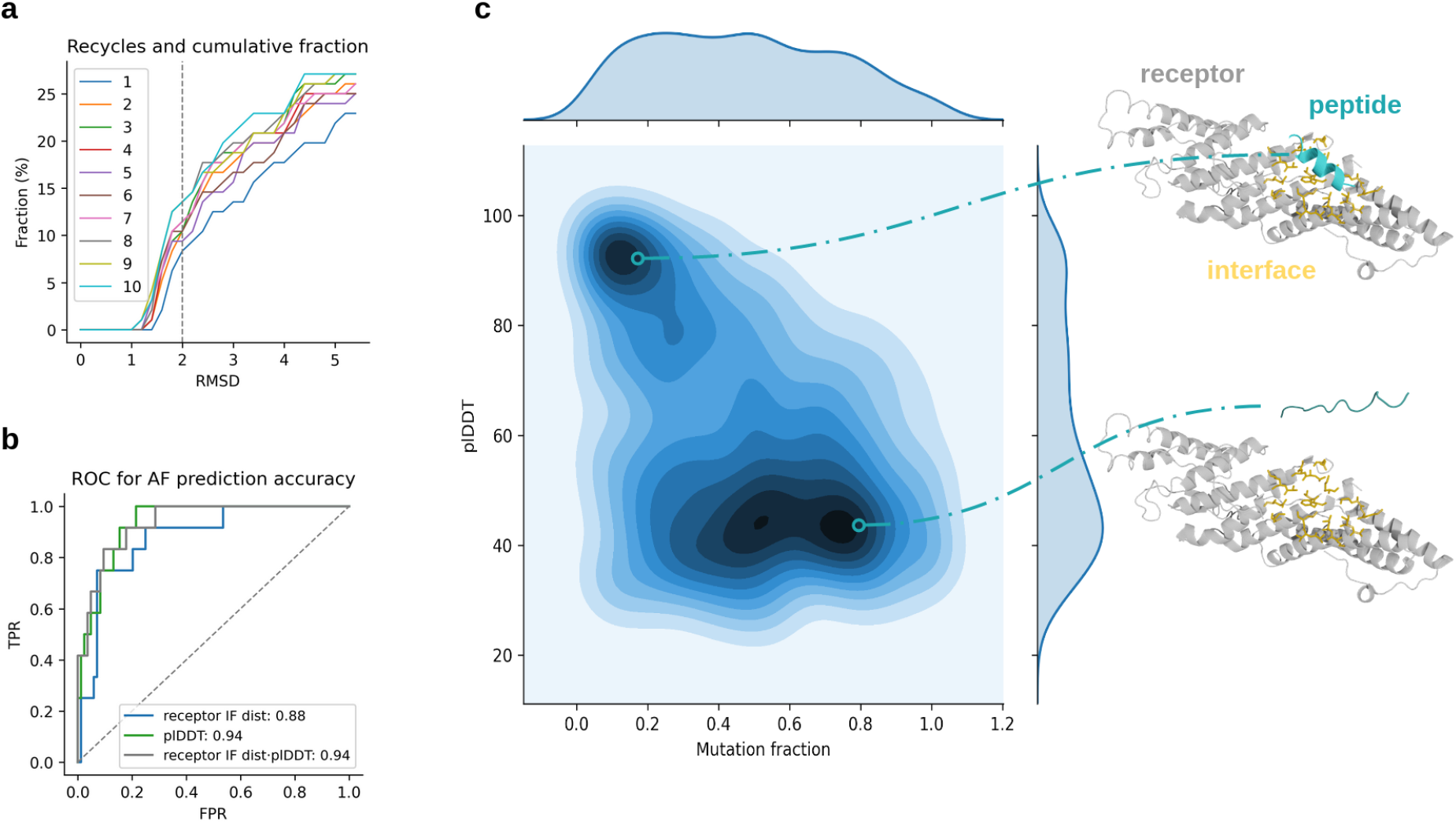
**a)** Analysis of the effect of the number of recycles on the outcome, measured by the average peptide interface (IF) RMSD. The IF is defined as Cβs within 8 Å between the peptide and its receptor protein, and the IF RMSD was calculated after aligning the CAs between the predicted and native receptor structures. The cumulative fraction vs the RMSD threshold (cut at 5 Å) is shown. At 8 recycles and above, no improvement at 2Å (grey dotted line) is found, although the difference is big between 1 and 8 recycles (12.5% vs 6.25%). **b)** ROC curve where positives (n=12; negatives, n=84) are predictions with an RMSD below 2 Å using 8 recycles from b). The plDDT of the peptide (plDDT) results in the highest AUC (0.94) and combining the plDDT with the distance from the receptor target residues to the peptide results in a slightly higher TPR at low FPR with no reduction in AUC. **c)** plDDT vs the fraction of mutated IF residues compared to the total peptide length using the peptides that could be predicted at 2 Å RMSD (n=12). In total, there are 1160 samples, 10 for each interface residue in the peptides. An illustrative example for PDB ID 3c3o (grey) is shown to provide intuition for this principle, where the peptide with a low mutation fraction (cyan, top) is highly ordered and close to the interface (orange) while the peptide with the high mutation fraction is disordered and further away (bottom).

Since not all peptide sequences can be predicted accurately at their true locations we analyse if we can distinguish when predictions are accurate. We analyse the predicted lDDT score (plDDT), a measure of how accurately each residue of a protein is predicted by AF and the distance to the target IF residues. The IF distance is calculated by taking the shortest distance from each atom in the target interface residues towards any atom in the peptide and then averaging. Figure 1b shows a ROC-curve where positives (n=12; negatives, n=84) have an average RMSD below 2 Å towards the true peptide structure IF. The plDDT of the peptide results in the highest AUC (0.94), and combining the plDDT with the receptor IF dist results in a slightly higher TPR at low FPR with no reduction in AUC. Together, this suggests that when AF is accurate in the prediction of protein-peptide interactions, the peptide will be situated close to the target interface and have high a plDDT score.

### Evaluation of AlphaFold specificity

AlphaFold may have learned certain binding pockets or potential interface regions, which would result in that any sequence could be predicted to interact with these regions. To analyse if it is possible to distinguish between potential peptide sequences to bind to a certain target interface, we mutate the 12 peptides from the set of 96 non-redundant peptides[10] that AF can predict at 2Å RMSD. We randomly alter amino acids in contact with the receptor in the native peptide sequences and predict the structure with AF (using 8 recycles). For each number of contact residues we introduce 10 new sequences, resulting in 10·L combinations for each peptide (1160 sequences in total for the 12 peptides with an average length of 12 residues).

The peptides with low mutation fraction have high plDDT (Figure 1c) with a strong concentration of samples at above 80 plDDT and below 20% mutations. The receptor IF distance tends to increase with the mutation fraction (Supplementary Figures 1 and 2) as well, and the peptides with low mutation fraction and low average distance to the receptor IF have high plDDT. It is also possible that the mutated peptides can still bind to the target interfaces, at least to some degree, which may explain why some mutated peptides are predicted close to the target interface. AF seems to be less sure of the placement of these sequences however, indicated by the lower plDDT (Supplementary Figure 1). By selecting peptide sequences with low average distance to specified target residues and high plDDT, it may be possible to retrieve true binders.

### *In silico* directed evolution

Since AF can distinguish between sequence changes in peptide binders using plDDT and the receptor IF distance, perhaps these metrics can be used to generate binding sequences. We create a directed evolution framework to test the possibility of designing peptide binders for certain interfaces using AF (Figure 2a). We input the length of the desired binder and a set of residues to target on the receptor. We predict the structure of the receptor-peptide complex using an MSA representation for the receptor and a single sequence for the peptide. Starting from a random sequence of length L, we mutate one amino acid randomly and keep the sequence for the next iteration if the receptor IF and the peptide IF distances are lower, the peptide centre of mass is closer to the target interface and the plDDT is higher than previously, as predicted by AF (*equation i*, Methods). We refer to this procedure as *in silico directed evolution*.

**Figure 2.**
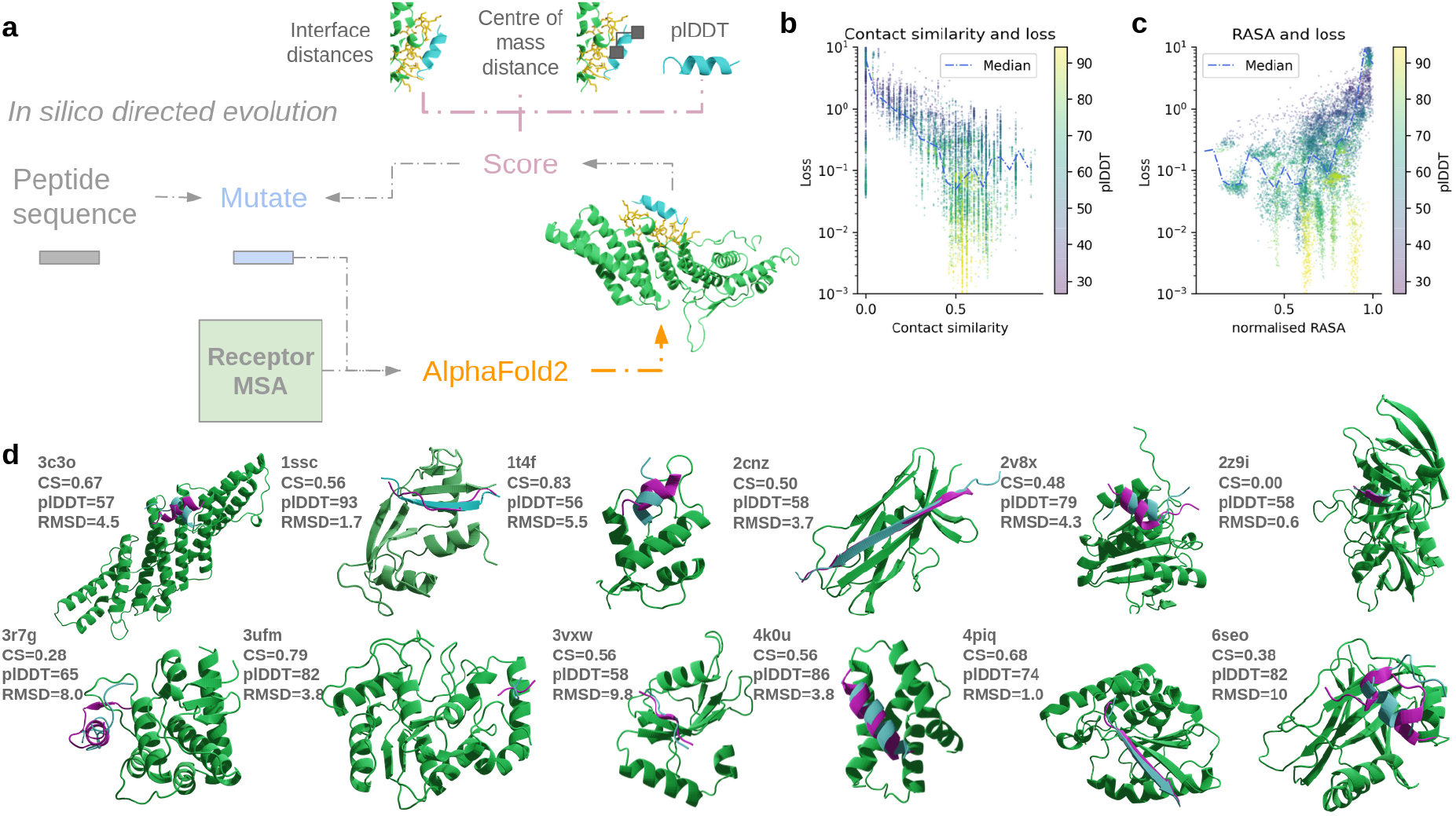
**a)** Depiction of the *in silico* directed evolution procedure. A random peptide sequence is input and mutated. The interaction between the peptide and a receptor protein is predicted with AlphaFold2 using a single sequence for the peptide, and an MSA for the receptor. An interaction score is calculated based on the distance from the peptide to the receptor target residues, a preferred centre of mass location (CA only) and the peptide plDDT. If the score is beneficial, the peptide sequence is kept for a new round of mutations. The mutation procedure is performed iteratively 1000 times. **b)** Contact similarity vs. loss during the optimisation procedure (12 peptides, 1000 iterations per peptide, n=12000). Each point is one sample and the line represents the median loss at different contact similarities (with step size 0.05). The loss is on a log scale. **c)** Relative accessible surface area (RASA) normalised by the highest value in each optimisation (corresponding to having the receptor residues fully exposed). Again, there is a strong relationship between the RASA and the loss function, although the lowest RASA values do not have the lowest losses. **d)** Final peptide designs from 1000 iterations (designed peptide: blue, native peptide: magenta, receptor: green) in superposition with the native structures (the receptor structure is only shown for the design). The median plDDT is 70 and median contact similarity (CS) 0.55. The PDB IDs for the upper and lower rows (left to right) are 3c3o, 1ssc, 1t4f, 2cnz, 2v8x, 2z9i and 3r7g, 3ufm, 3vxw, 4k0u, 4piq, 6seo, respectively.

The peptide IF distance is the average of the shortest distance from all atoms in the peptide to any atom in the target residues. This metric was included to ensure that all residues in the peptide will be close to the target residues during the optimisation. We noticed that using only the receptor IF distance, can result in parts of the designed peptides not interacting with the target residues for longer sequences (Supplementary Figure 3). The peptide centre of mass (using only CAs) was added to ensure that a peptide binding to the desired interface of the target residues is created. We noticed that for some proteins, it is possible to obtain designs towards opposite sides of target residues with similar distances (Supplementary Figure 4). The *in silico* directed evolution was performed for the 12 peptides from the set of 96 non-redundant peptides[10] that AF can predict at 2Å RMSD. The optimisation was run for 1000 iterations per peptide, using 8 recycles per iteration (see Supplementary Figure 5 for the optimisation curves).

### Performance analysis of the *in silico* directed evolution

To analyse how well the *in silico* directed evolution works, we analyse the similarity between the residues interacting with the receptor interface in the native and designed structures (contact similarity, see methods) and the relative accessible surface area (RASA) of each residue in the receptor interface. Figure 2b displays the relationship between the contact similarity and the optimisation loss coloured by plDDT, showing an increasing contact similarity with a decreasing loss. A saturation at around 50% contact similarity is found, which is also where the highest plDDT scores are found. The median contact similarity is 56%, taking the models with the lowest loss from the optimisation and the median plDDT is 70. The RASA increases with the loss, suggesting that sequences likely to be energetically favourable for binding are generated (Figure 2c, median=8% unnormalised). The loss converges before 500 iterations for all optimisations (Supplementary Figure 5).

Normally the sequence identity is analysed for protein design applications of entire proteins. For short sequences such as peptides, this is a relatively bad measure as many different peptides may in fact bind to a certain interface and it is not known how the sequence relates to the structure for protein-peptide interactions. Several different peptide residues interact with the same receptor residues as well (Supplementary Figure 6), suggesting that the exact order of some residues may not be that important. Figure 2d shows the designs with the best loss from the optimisation in superposition with the native structures. In many cases, the designs are very similar to the native structures (median CA RMSD=4.3 Å). However, it is possible to obtain a different structure with a high plDDT (e.g. 6seo, plDDT=82, RMSD=10) or a very similar structure with a relatively low plDDT (2z9i, plDDT=58, RMSD=0.6 Å). The reason not all designs have high plDDT (>80) may be due to the optimal solution not being reached. We explore this further in the convergence analysis section below.

### Structural flexibility: the receptor structure changes to accomodate the binder

The sequences of the designs are different from the native ones, even though the median contact similarity is 56%. To accommodate different peptide sequences, it is likely that the structure of the receptor interface also has to change. When analysing the RMSD of the receptor interface Cβs vs. the contact similarity (Figure 3a) compared to the native structures during the optimisation, one can see that above 1 Å RMSD, the contact similarity decreases rapidly. This suggests that AF does take the interaction with the peptide structure into account when predicting the structure of the interface with the receptor. Most structures are predicted to have approximately 0.5 contact similarity and these have the highest resemblance with the native interface as well (below 0.5 Å Cβ RMSD).

**Figure 3.**
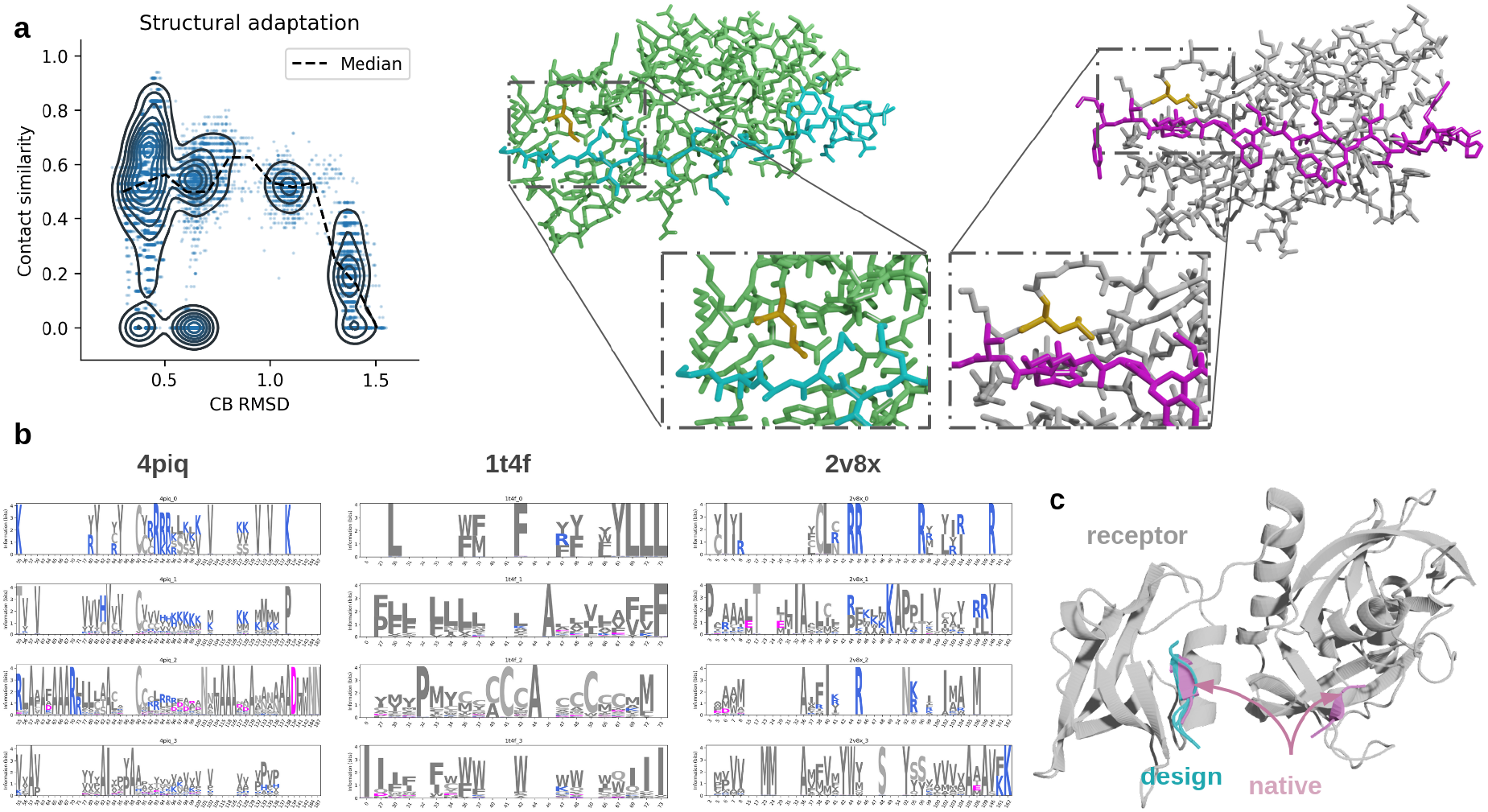
**a)** Analysis of the structural flexibility during the optimisation for the peptides that could be predicted at 2 Å RMSD (n=12). RMSD of the receptor Cβs vs the contact similarity. Each point represents one iteration (n=12000) and the line the running median with a step size of 0.1 RMSD. KDE curves are also included to show the data distribution. At low RMSDs, the contact similarity is high suggesting that the receptor structure adapts to the peptide structure. An example for PDB ID 2cnz is shown, where residue 12 (orange) in the receptor (grey, green) changes orientation to interact with the peptide residues (blue, magenta). **b)** Convergence analysis. Contact sequence logos using the top 10% designs from three runs for PDB IDs 4piq, 1t4f and 2v8x (see Supplementary Figure 6 for all 12 peptides). The amino acids are coloured by type. c) Analysis of alternative binding site retrieval. Optimising only for the plDDT using 500 iterations and three different initialisations for PDB ID 2z9i. The peptides with the highest plDDT scores were selected. Only one of the two possible binding sites (native, magenta) on the receptor protein (grey) are recovered by the designs (blue).

### Convergence analysis

To analyse if the sequence converges in the *in silico* directed evolution procedure, we initialise the procedure three times and compare the outcome. We calculate the frequencies of any amino acid in the peptide being in contact (Cβs within 8 Å) with any amino acid in the receptor. From these frequencies, we create “contact logos”, representing the likelihood of certain amino acids between the peptide and receptor being in contact throughout the optimisation. Figure 3b shows the contact logos from each of the three optimisation runs compared to the contacts in the native structure using the top 100 designs (10 %) for PDB IDs 4piq, 1t4f and 2v8x (see supplementary Figure 6 for all 12 peptides). Each position in the logo represents a position in the receptor interface and the amino acids the probabilities of having a contact in the peptide to that residue.

The designs do not converge overall, although several receptor-peptide contacts do. For 4piq, C is kept in the first two runs followed by a charged region. In position 55, there is a K in the native vs. R in run 2. In positions 127,128 there are K,K in run 1 and VV in run 3 vs. K,K and V,V in the native sequence. Run 3 also has 132,135 V,V identical to the native sequence. For 1t4f, there are almost only polar residues. In position 30 there is a L in the native and run 1 and in 36,37, WF, FM compared to WF in run 3. All runs of 2v8x have A in pos 36, but this is absent from the native sequence. The native 2v8x has R in 44,45 also present in run 1 (44) and 2 (45), but not in run 3. Run 1 has 38,40 LC and the native C,L. Likely these can be switched equivalently displaying again the possible variation of positions in 3D.

### Finding alternative binding sites

Some receptors have alternative binding sites, bound by different peptides. It is possible that by not specifying the target interface residues and centre of mass, peptides binding different interfaces (or none) are generated. To analyse this, we use the receptor from PDB ID 2z9i which is reported to bind two different peptides (4 and 5 residues long, respectively, Figure 3c). Optimising only for plDDT and a length of 5 residues, we take 500 steps mutating one amino acid at random and run three different initialisations. We select the complexes which have the highest plDDTs from each initialisation and analyse the variation in binding site.

There is no variation in the resulting binding sites, suggesting that some sites are easier to optimise towards. Therefore, an approach using only the plDDT as a loss function to design a binder towards a specific site or to extract different binding sites seems unlikely to be successful. However, this approach may be used to find the most promising binding sites of a target protein, although it is unknown if these will be relevant for modulating protein functions.

### Distinguishing miniprotein binders

Miniproteins are small protein scaffolds with sizes only slightly above the general peptide limit (50 residues). Such proteins have been designed to bind target interfaces, with limited success[3]. To see if the modification of the AF protocol described here can distinguish these binders we analyse sequences tested against four different receptor proteins with solved receptor-binder structures (FGFR2: https://www.rcsb.org/structure/7N1J, TrkA: https://www.rcsb.org/3d-view/7N3T/1, IL7Ra: https://www.rcsb.org/structure/7OPB and VirB8:https://www.rcsb.org/structure/7SH3) [3]. These binders were evaluated by counting the number of times they were detected to interact on yeast cell surfaces by Fluorescence-activated Cell Sorting (FACS) and Next-Generation Sequencing (NGS counts). We sampled 1000 sequences below 1000 NGS counts (to reduce the computational cost, 5578 sequences in total) and all above and predicted the receptor-miniprotein structures with AF using the same protocol as described above.

Figure 4 displays the relationship between the loss function used for the in silico directed evolution and the normalised NGS counts. At NGS counts of zero, there is a much higher tendency to obtain a high loss than compared to at 0.04 where the loss is close to 0. As a result, 47% of binders can be selected at a FPR of 20% (Supplementary Figure 8). Compared to the mutated peptide binders, the average plDDT is much higher (84 vs 58) suggesting that AF is unsure about the residue locations of unbound peptides but not of unbound miniproteins. This corresponds well to the higher flexibility of peptides which likely only achieve their native configuration upon binding, reflected in the ability of AF to account for structural flexibility upon binding (Figure 3a).

**Figure 4.**
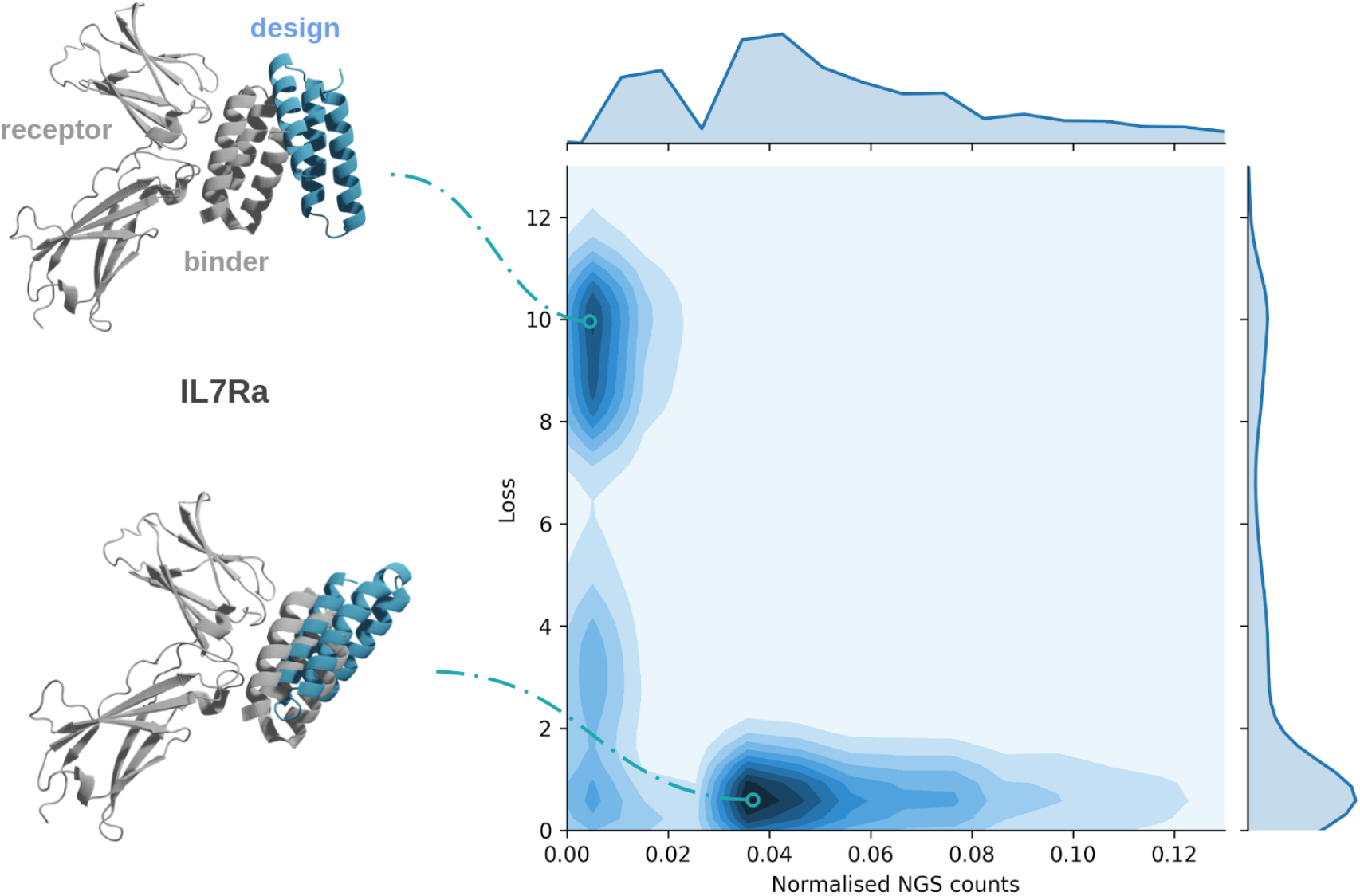
Loss and normalised NGS counts for the four tested miniprotein binders with resolved structures (n=5578 of which 2782 have NGS count 0). An illustrative example at high and low loss for IL7Ra is shown in relation to the resolved structures (native grey, design blue). The NGS counts are normalised by dividing with the highest count. The axis limits have been cut at 13 and 0.13 for the loss and normalised counts, respectively. For the full range, see Supplementary Figure 8 and for a ROC curve for selecting binders using the loss, see Supplementary Figure 9.

### Peptide binding affinity

So far, we have only assessed the ability of AF to distinguish known peptide-receptor structures and mutated versions of these in a qualitative manner (binding/not). It is also desirable to assess affinity differences between peptides in a real setting. To analyse if AF can distinguish between peptide binding affinities we analyse the ELISA ratios of a set of 234 peptides towards 9 receptor proteins from different peptide recognition modules (PRMs) from http://www.prm-db.org/; [20]. The highest affinity binders are those with the highest ELISA ratio values. As can be seen in Figure 5a, there is a weak relationship between the ELISA ratio and the plDDT. At the highest densities, the ELISA ratios are approximately 13 and 23 vs 35 and 55 for the plDDT. The plDDT values are quite low overall, suggesting that AF is not very confident in the predictions.

**Figure 5.**
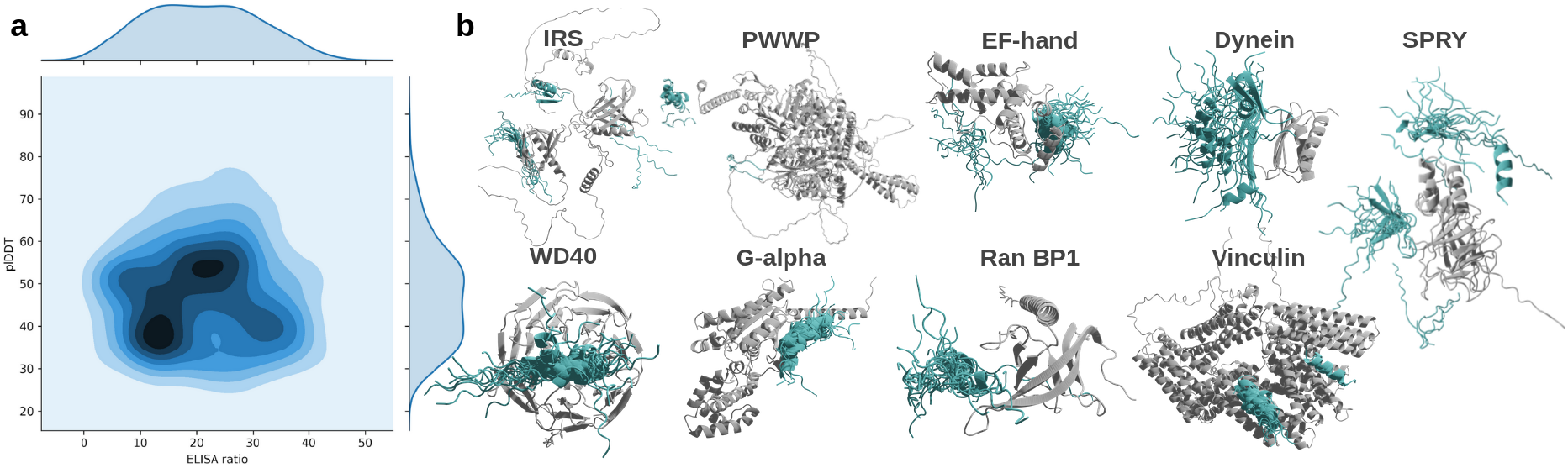
Analysis of peptide binding affinity. **a)** Density plot (n=234) of the plDDT of the peptide vs the ELISA ratio. There is a weak relationship between ELISA ratio and the plDDT and the number of contacts from the AF predictions. **b)** Structural superposition of all predictions for each PRM with the receptor structures in grey and the peptides in blue. When analysing the predictions, one can see that most of them display variation in binding sites. The UNIPROT IDs for the PRMs are Q99704, P52701, O75340, P63167, Q99619 P61964, P19086, P43487, P18206 (left to right, respectively).

When analysing the predictions, one can see that most of them display variation in binding sites (Figure 5b). This may partially explain why the ELISA ratios are difficult to distinguish. Ratios across different binding sites may not be comparable due to some binding sites being weaker than others and still produce strong confidence in the plDDT and number of contacts. To properly evaluate this idea, one would have to know the true binding sites of these peptides and the exact contacts. Affinity measurements can however not distinguish between binding sites, only report binding. Regardless, the plDDT values of the predicted peptides are too low to distinguish between true binders according to Figure 1c.

### Binding motif recovery

Motif-driven receptor-peptide interactions have proven to be easier to predict with AF than the set of peptides analysed here[10]. There may be less variation in the possibility to bind a certain interface region when there are strong motifs. This suggests that it may also be possible to recover binding motifs through the *in silico* directed evolution procedure. Using a set of 12 protein-peptide complexes with known motifs, and structures [10] [21], 1000 iterations of binding optimisation were performed. Figure 6 shows sequence logos, representing the frequency of amino acids for each position in the receptor interfaces that interact with a motif amino acid in the native peptides created with the top 100 designs (10%) for each peptide.

**Figure 6.**
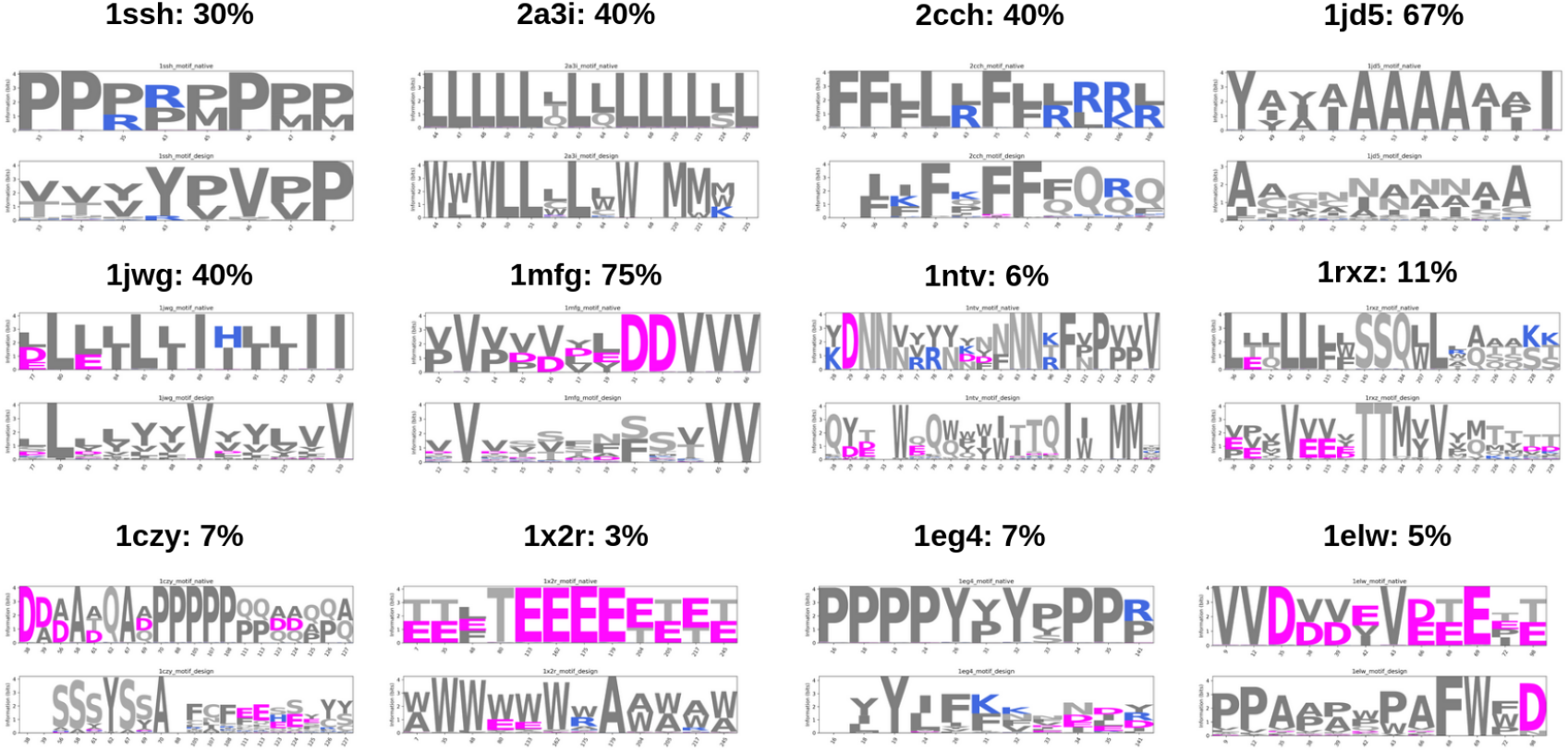
Analysis of the recovery of binding motifs. Amino acid frequency logos using the top 100 designs (10%) for each peptide (top row=native sequence, bottom row=design sequences). The logos represent the information at each position of the receptor interfaces that interact with a motif amino acid in the native peptides. The amino acids are coloured by type and the motif similarity (*equation iii*, Methods) between the native and design amino acid frequencies are displayed as fractions. The matching motif positions for each PDB id are: 1ssh: RP-PP pos 43-48, 2a3i: L-LLLL pos 47-64, 2cch: FF-(K)FF–R pos 36-106, 1jd5: [A/I][I/A][A/I]AAAA[A/I]A pos 49-66, 1jwg: [L/D]LLLLL–LL pos 77-125, 1mfg: VVVVVD(D)--VVV pos 12-66, 1ntv: D pos 29, 1rxz: EQ—---Q—QTT-[K/T] pos 40-229, 1czy: no matches, 1×2r: E pos 133, 1eg4: R pos 141, 1elw: P(D) pos 72-98.

Motifs are linear due to sequence analysis reasons. In reality, many different rearrangements of the same motif can likely bind well to a target region in 3D. Analysing linear motifs when assessing binding in 3D is thereby a flawed concept, which is why we instead analyse interactions between motif and receptor residues. The motif similarity is calculated by comparing the native and design amino acid frequencies (*equation iii*, Methods). The best reproduced motifs are 1jd5 and 1mfg which correspond very well to the native receptor contacts with 67 and 75% motif similarities, respectively, and the worst 1×23 and 1elw with 3 and 5% motif similarities, respectively. As the structure changes to accomodate binding (Figure 3a), it is possible that many different amino acids of the same class can bind the same interface residues which may explain some of the low motif similarities.

### Conclusions and Future outlook

We have demonstrated that AF can distinguish between mutated and native peptide binders, recover amino acid contacts and reduce receptor surface accessibility using the *in silico* directed evolution procedure. Peptide binding motifs can be recovered, a meaningful application on its own, and designed minibinders can be distinguished from non-binders. It is not possible to predict the affinity of binders, possibly due to AF being unsure about these predictions reflected by the low plDDT values.

The *in silico* directed evolution platform is not proposed as to create final drug designs, but as an initial starting point for further laboratory analysis and optimisation. Narrowing the vast search space of possible sequences as described here enables reducing costs as well as developing timelines for new binders substantially. Since no structural scaffold is required, AF is free to adapt binders to any conformation it sees fit, allowing for structural flexibility of the receptor interface.

Although several additional aspects such as solubility, immune system interactions and aggregation propensity have to be considered, perhaps we are now one step closer to being able to tackle diseases with targets suitable for peptide drugs that are not considered financially viable today. Likely, such aspects will also be possible to consider in future computational frameworks, ultimately leading to very few laboratory tests being necessary. We do however not expect to be able to predict peptide binders for any target. AlphaFold does indeed fail for 84 of the 96 analysed peptides at a 2Å RMSD threshold. Still, we do expect that our computational framework can aid experimentalists substantially. As we have shown here; if AlphaFold can predict a peptide binder close to its native conformation, the plDDT score is likely to be high and we can then design a sequence that is likely to bind to the target interface.

## Methods

### Peptide structural data

The peptide dataset was taken from a recent study[10] where 96 non-redundant protein-peptide structures were extracted from the PDB and manually analysed to ensure structural divergence (each involves a distinct fold). This dataset was created as follows quoting the original publication[10]:

- *The PDB was queried for entries with two chains only, and filtered for those having possible protein-peptide interactions according to the following criteria:*
  - *1. One chain must be over 30 amino acids long, and one chain must contain between four and 25 amino acids (with at least three amino acids resolved in the solved structure)*.
  - *2. The peptide chain must have at least two residues within 4 Å distance from the protein chain. This yielded a total of 16,931 structures belonging to 1102 ECOD domains*^52^.
- *Once possible interactions were identified, the following filters were applied:*
  - *1. Remove structures with peptide residues annotated as UNK*,
  - *2. PDB-range and seq-range fields must agree on the indices of the receptor domain according to ECOD annotation*,
  - *3. Apply symmetry operations (from the PDB entry) on the asymmetric unit and check for possible crystal contacts that may affect the bound conformation of the peptide*.
- *Remove cases where at least 20% of the peptide residues are in contact with symmetry mates*.
- *Structures from ECOD families represented in the motif and non-motif sets were removed. The resulting list was manually validated, and structures were set aside that contain a peptide conformation that might be influenced by context not included in the input (e*.*g*., *structures containing ligands in the vicinity of the peptide binding site or peptides with modified residues, such as PTMs)*.
- *The final list after filtering and manual validation consists of 96 peptide–protein complexes*.*”*

### Structure prediction with AlphaFold2

A modification of AlphaFold (v2.0) [22] (AF) based on the FoldDock protocol[4] and a recent study for protein-peptide structure prediction[10] was run, where the receptor is represented as an MSA and the peptide as a single sequence. The MSA was constructed from a single search with HHblits[23] version 3.1.0 against uniclust30_2018_08[24] using the options:

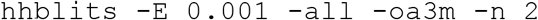

The MSA of the receptor was input together with the single sequence representation of the peptide to the AF folding pipeline, using model_1, one ensemble and between 1-10 recycles (8 were found to be optimal, see Figure 1a). In model_1, no predicted TM-score (pTM) or frame aligned point error (FAPE) is available, only the predicted lDDT (plDDT) for each residue.

The structural prediction was performed on one NVIDIA A100 Tensor Core GPUs with 40 Gb of RAM. On average, compiling the folding pipeline took 144 s and each iteration took 46 s, resulting in an average total time of 144+46·1000 ≈ 12 hours and 49 minutes for 1000 iterations.

### Sequence optimisation

To optimise the peptide binding sequence, one amino acid was changed at each iteration starting from a random sequence initialised with a Gumbel distribution over all standard 20 amino acids. The change was conditioned on not resulting in any previously evaluated sequence. This change (or mutation) was evaluated using AF by predicting the interaction with the receptor structure and calculating a loss defined as:

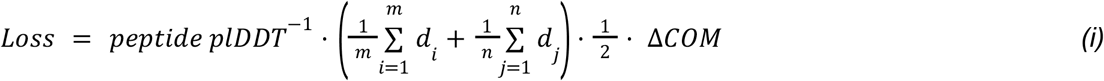

Where the peptide plDDT is the average plDDT over the peptide, di is the shortest distance between all atoms in the receptor target atoms and any atom in the peptide, dj is the shortest distance between all atoms in the peptide and any atom in the receptor target atoms and ΔCOM is the CA centre of mass distance between the native and predicted peptides.

### Contact similarity

To analyse how similar the designed peptides are to the native ones, we compare the contacts (defined as beta carbons (Cβs) within 8 Å from each other) between the receptor and the native peptide and of the receptor and the design. We group amino acids into five different categories (see below) depending on their physical characteristics to capture the fact that many different amino acids may interact with receptor interface residues equally well (e.g. different positive residues). From these groupings, we calculate the fraction of interactions in the native peptide that is preserved in the design, which we call contact similarity.

For each position in the native receptor interface, the interacting residues are extracted and the unique groupings are annotated. This means that repeat contacts are not counted, e.g. if residue one interacts with A,F and R, this translates to Hydrophobic and Positive. The two counts of hydrophobic residues (A and F) are not considered. This is because contacts may vary and it may be sufficient to have one stronger hydrophobic interaction as compared to two weaker ones.

#### Amino acid categories

Hydrophobic: A, F, I, L, M, P, V, W, Y

Small: G

Polar: N, C, Q, S, T

Positive: R, H, K

Negative: D, E

### Relative accessible surface area

To calculate the relative accessible surface area (RASA), we use DSSP (version 3.0.0). For each residue we extract the area and normalise it according to empirical measurements[25], with a maximum allowed value of 100 %). Polar, positive and negative residues were given RASA values of 0 as the exposure of these can be energetically favourable. To compare RASA values between different peptides, we normalise them with the maximum observed values during optimisation corresponding to exposed surfaces.

### Miniprotein binders

Four different designed miniprotein binders that had resolved structures towards single-chain proteins were selected from a recent study (FGFR2: https://www.rcsb.org/structure/7N1J, TrkA: https://www.rcsb.org/3d-view/7N3T/1, IL7Ra: https://www.rcsb.org/structure/7OPB and VirB8:https://www.rcsb.org/structure/7SH3) [3]. These binders were evaluated by counting the number of times they were detected to interact on yeast cell surfaces by Next-Generation Sequencing (NGS counts) in subsequent pools obtained from Fluorescence-activated Cell Sorting (FACS). We used the final pools for each of the four receptors as the designs there will represent the strongest binders. In total there were 172581 sequences available in this study and to limit the computational requirements we sampled up to 1000 sequences below 1000 NGS counts and all above and predicted the receptor-miniprotein structures with AF as described above. In total, 5578 receptor-miniprotein structures were evaluated, 2013 for FGFR2, 1099 for TrkA, 1249 for IL7Ra and 1217 for VirB8 (see Supplementary Figure 7 for the NGS count distributions).

### Peptide affinity analysis

To analyse if AF can distinguish the affinity of peptide binders, we selected 255 peptides from 10 different receptor proteins from different peptide recognition modules (PRMs) that have reported ELISA ratios using the PRM-db[20]. We selected the targets at random (the first that came up in the search). The full length uniprot sequences were extracted and the same prediction procedure as above (hhblits MSA representation for the receptor and a single sequence for the peptides, 8 recycles) was performed to predict the structures. One of the selected targets (PABP, 2799 residues) was too large to fit in memory resulting in 21 OOM errors and a total of 234 peptide binder predictions for 9 different receptor proteins.

### Amino acid frequency logos

To create amino acid frequency logos, the number of each type of amino acid was counted at each position (i.e. receptor interface position) and normalised with the total number of counts at that position. The logos were created using the python package Logomaker [26]. We added a pseudo probability of 0.001 to the counts to handle otherwise infinite negative logarithms. All logos are made using bit information (Shannon entropy[27]) defined as:

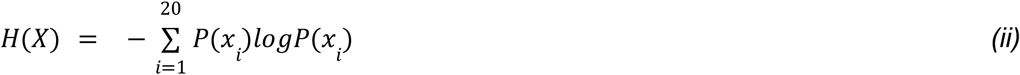

Where P(x) is the probability (or frequency) to observe amino acid i in a certain position and H(X) is the total bit information. Logos from the native AA sequences were created for comparison.

The similarity between two amino acid logos are calculated by multiplying the frequency of each amino acid at each position and averaging over all positions accordingly[28]:

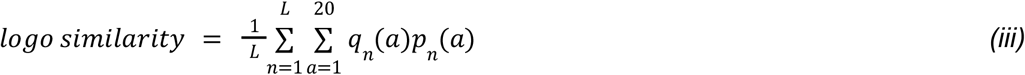

Where L is the number of positions in the logo and q_n_(a) is the frequency of amino acid a in position n for logo q and p_n_(a) that for logo p. The similarities were normalised with the comparison of the native logo to itself to have an equal scaling between zero and one.

### Binding motif recovery

To analyse if it is possible to recover binding motifs using the *in silico* directed evolution procedure, 1000 iterations of optimisation (using the same settings as above) was run for a set of 12 protein-peptide complexes with known motifs and structures [10] [21]. The receptor interface positions that bind a motif residue in the native structures were extracted and compared to the top 100 designs (10 %) from the optimisation. Amino acid frequency logos were created by counting what amino acids in the peptides interact with certain receptor interface positions and thereafter calculating the bit information (above). All contacts are defined as having <8 Å between Cβs.

## Acknowledgements

All protein structures were visualised using blender (https://www.blender.org/). Financial support: Swedish Research Council for Natural Science, grant No. VR-2016-06301 and Swedish E-science Research Centre and from Knut and Alice Wallenberg foundation. Computational resources: Swedish National Infrastructure for Computing, grants: SNIC 2021/5-297, SNIC 2021/6-197, Berzelius-2021-29 and Berzelius-2022-106. A.E received all financial support and computational resources.

## Author contributions

PB designed and performed the studies and analyses. PB wrote the first draft of the manuscript and prepared all figures which were later edited and improved by AE and PB. AE obtained funding.

## Availability

All results and information required to reproduce this study are available from: https://gitlab.com/patrickbryant1/binder_design. For local installation of EvoBind see: https://github.com/patrickbryant1/EvoBind. A web version through Google colab is available here: https://colab.research.google.com/github/patrickbryant1/EvoBind/blob/master/EvoBind.ipynb.

## Supplementary material

**Supplementary Figure 1.**
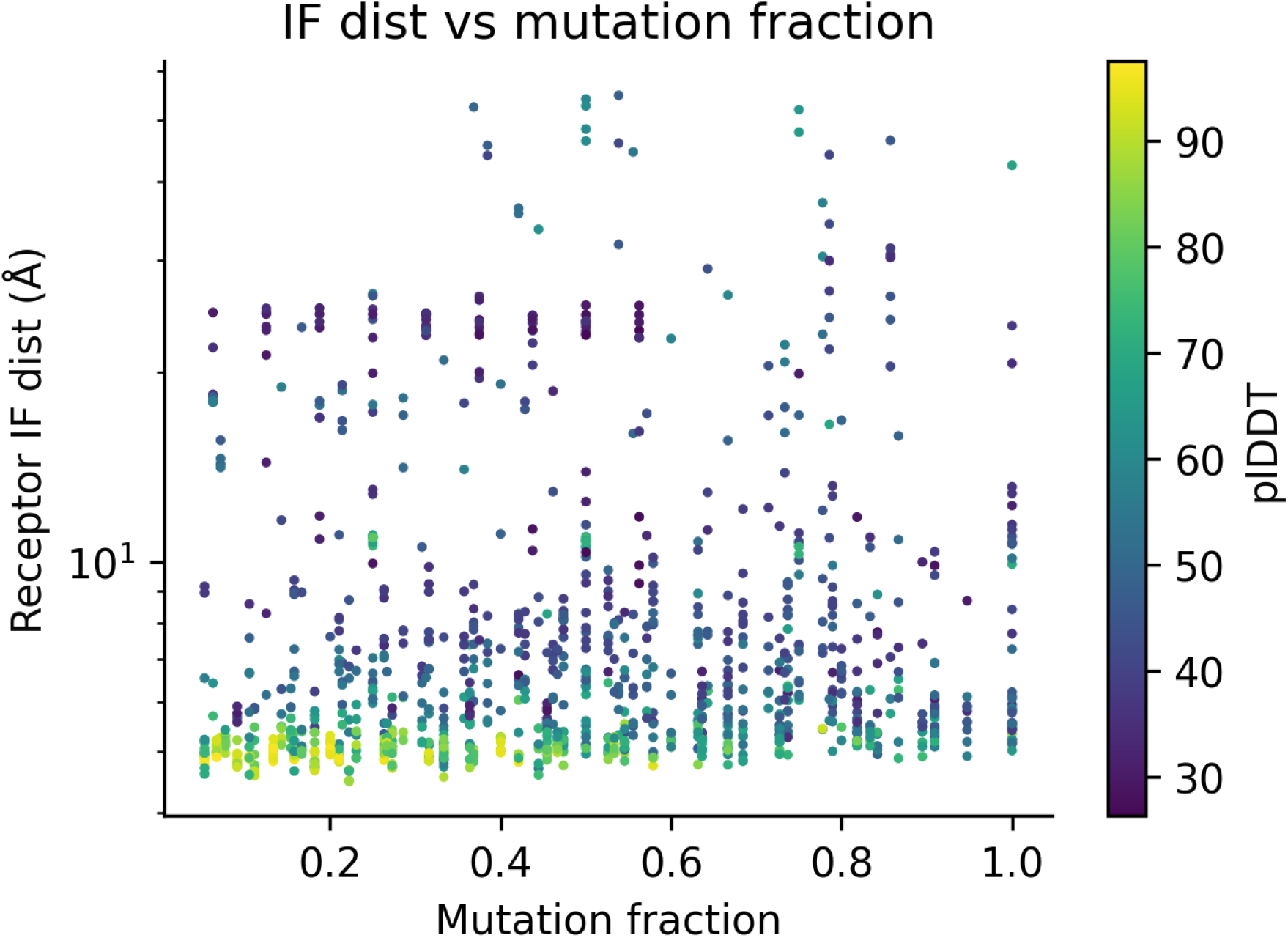
Relationship between the mutation fraction and distance to the receptor interface for the 12 peptides that could be predicted using AF at <2Å RMSD. Each sample (n=1160) is colored by plDDT. The y-axis starts at 4 Å and is displayed in logarithmic scale.

**Supplementary Figure 2.**
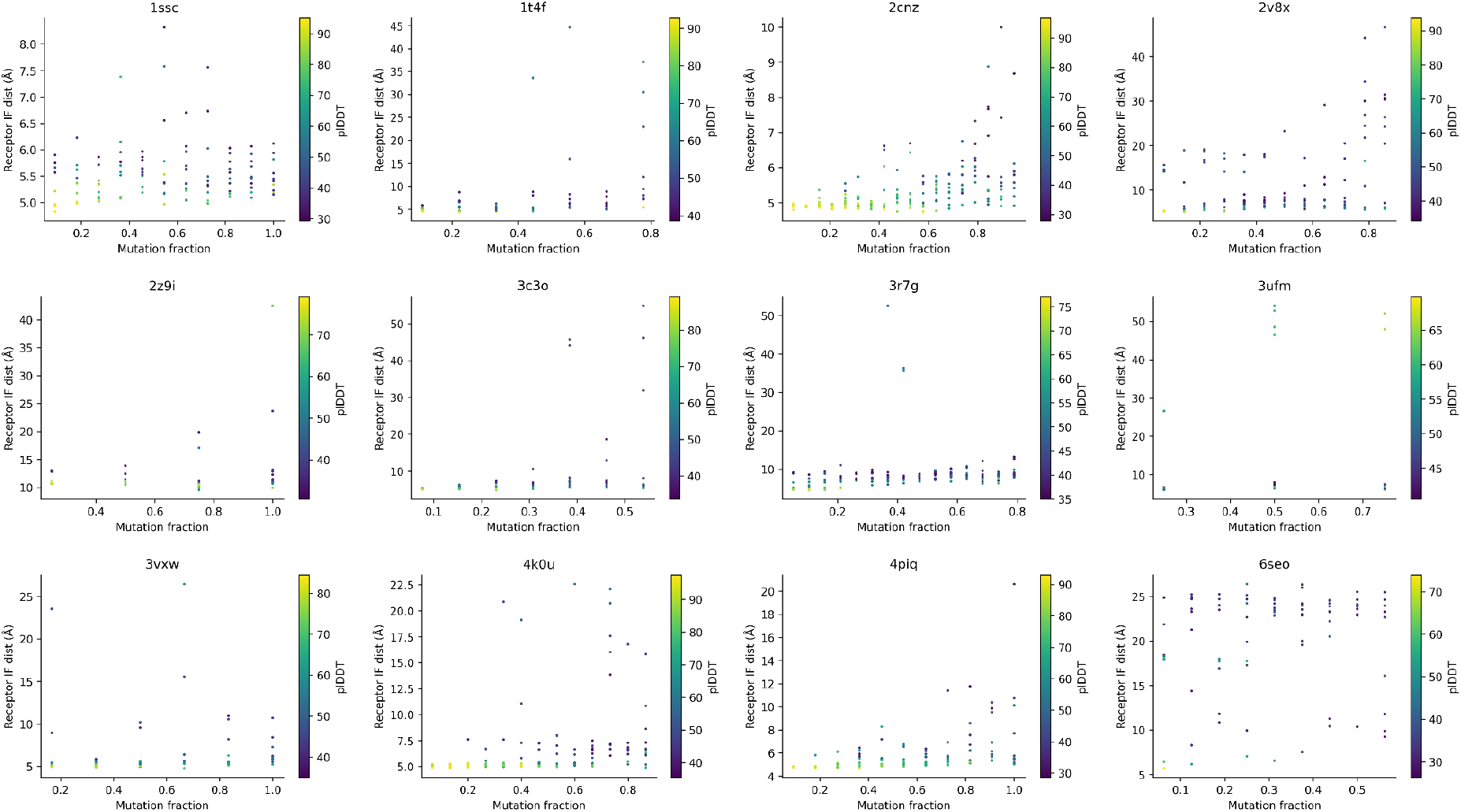
Relationship between the mutation fraction and distance to the receptor interface for each of the 12 peptides that could be predicted using AF at <2Å RMSD. Each sample is colored by plDDT.

**Supplementary Figure 3.**
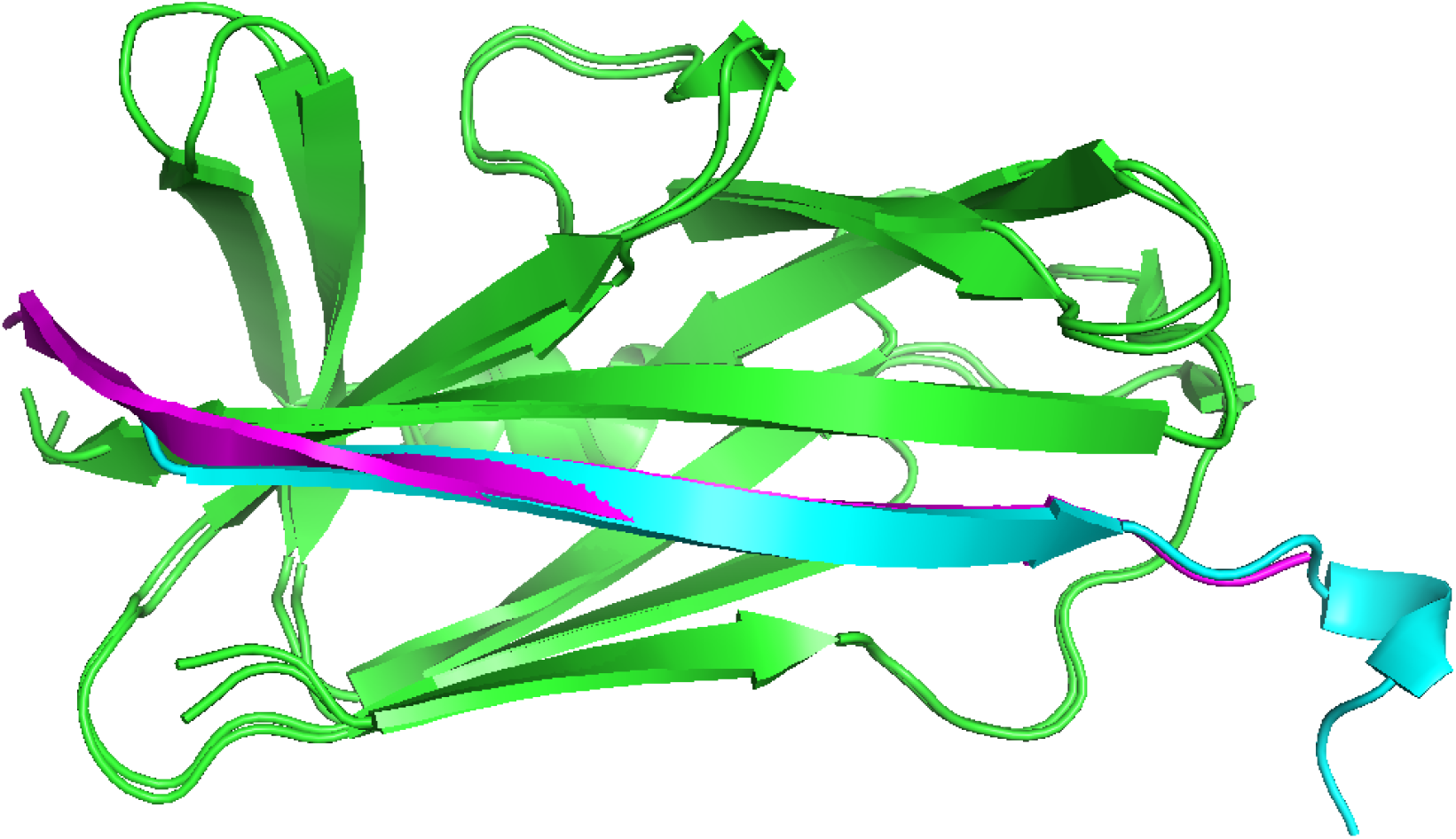
Best model obtained from the 2cnz optimisation using only the Receptor IF distance and the plDDT. The receptor structure is shown in green and the designed and native peptides in blue and magenta, respectively. One part of the designed peptide protrudes and does not interact with the target interface.

**Supplementary Figure 4.**
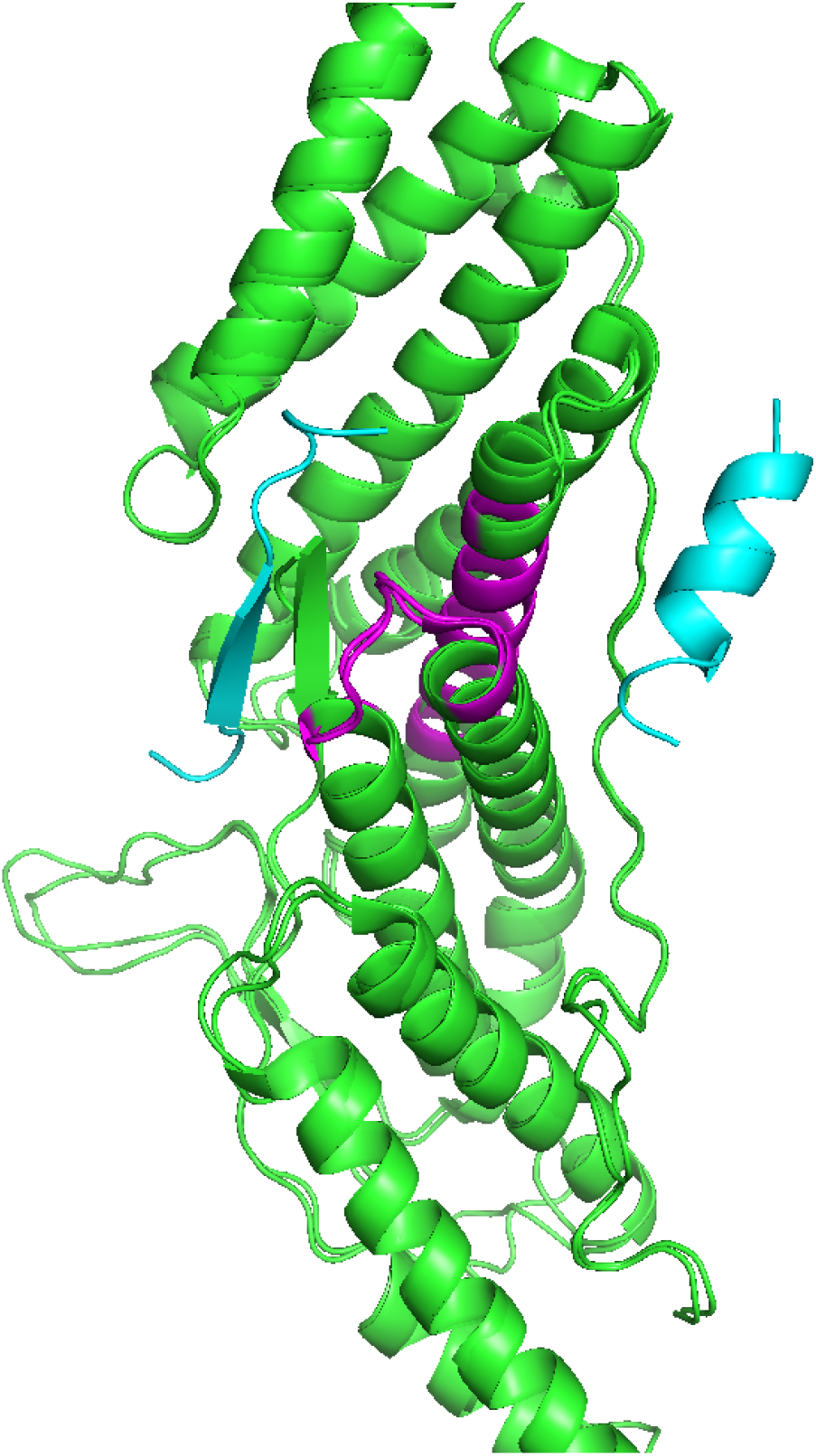
Without using the peptide centre of mass in the design procedure, it is possible to design a peptide binding to the wrong side of the target residues. An example for 3c3o is shown here, where the design binds the wrong side of the target (magenta=target, blue left=design, blue right=native).

**Supplementary Figure 5.**
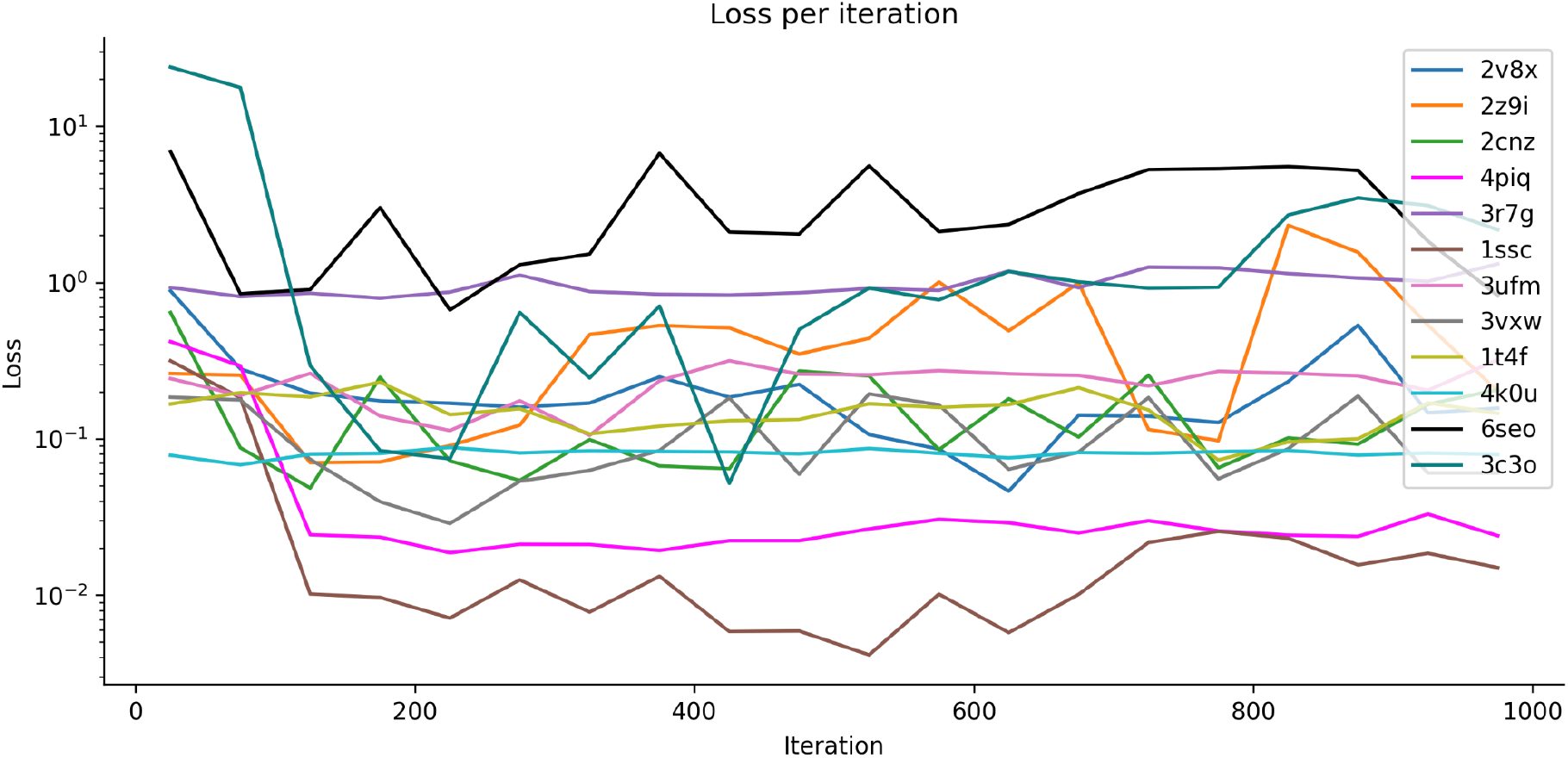
Median loss per iteration calculated using a step size of 50 iterations for each of the 12 peptides that could be predicted using AF at <2Å RMSD. The loss converges at around 200 iterations for most peptides. Note the logarithmic scale on the y-axis.

**Supplementary Figure 6.**
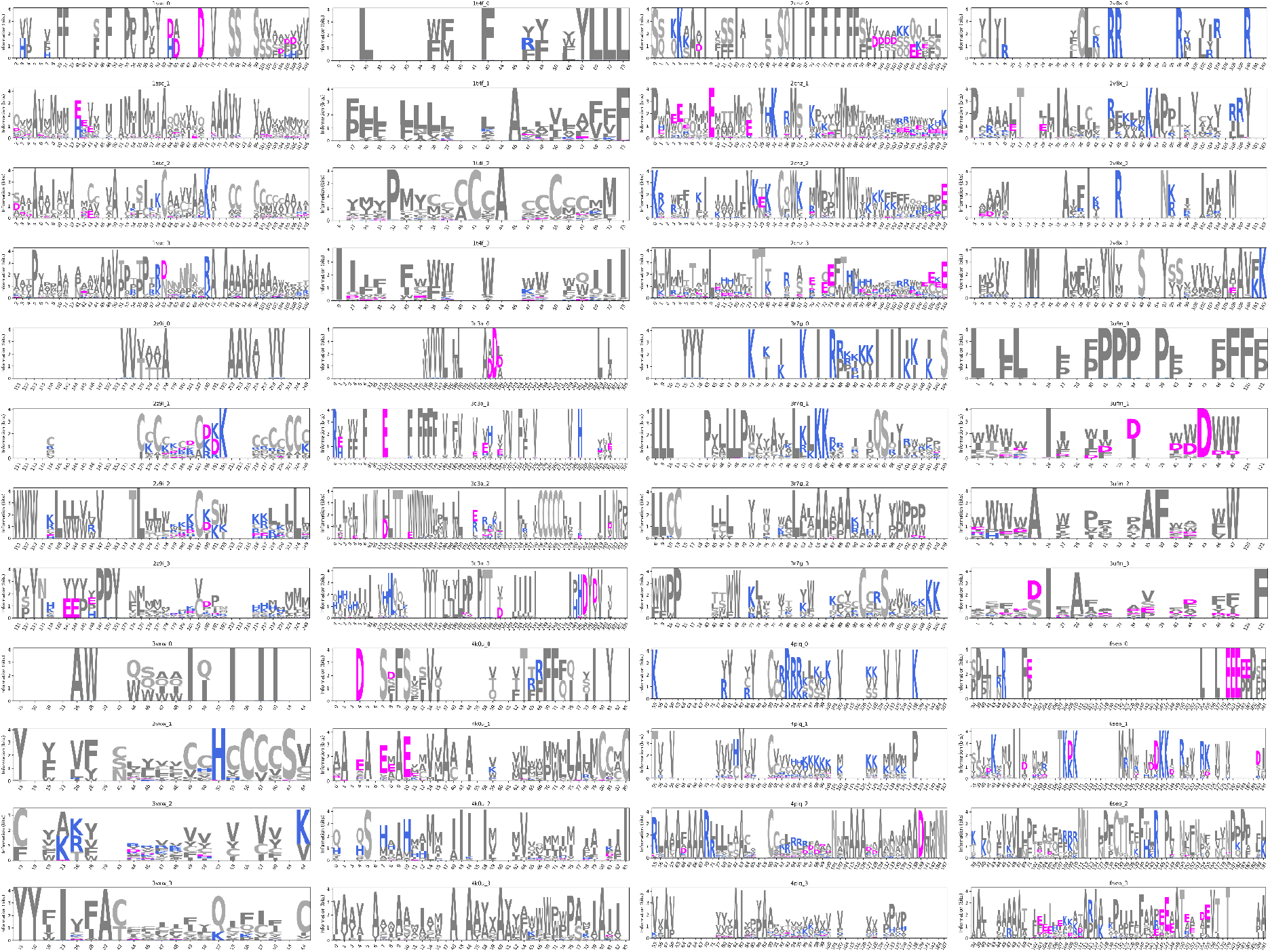
Convergence analysis. Contact logos for each position in the receptor interface in contact with any amino acid in a peptide using the top 10% (100) designs for three different runs. The top logo for each PDB ID represents the native sequence and below are the logos resulting from the three different runs.

**Supplementary Figure 7.**
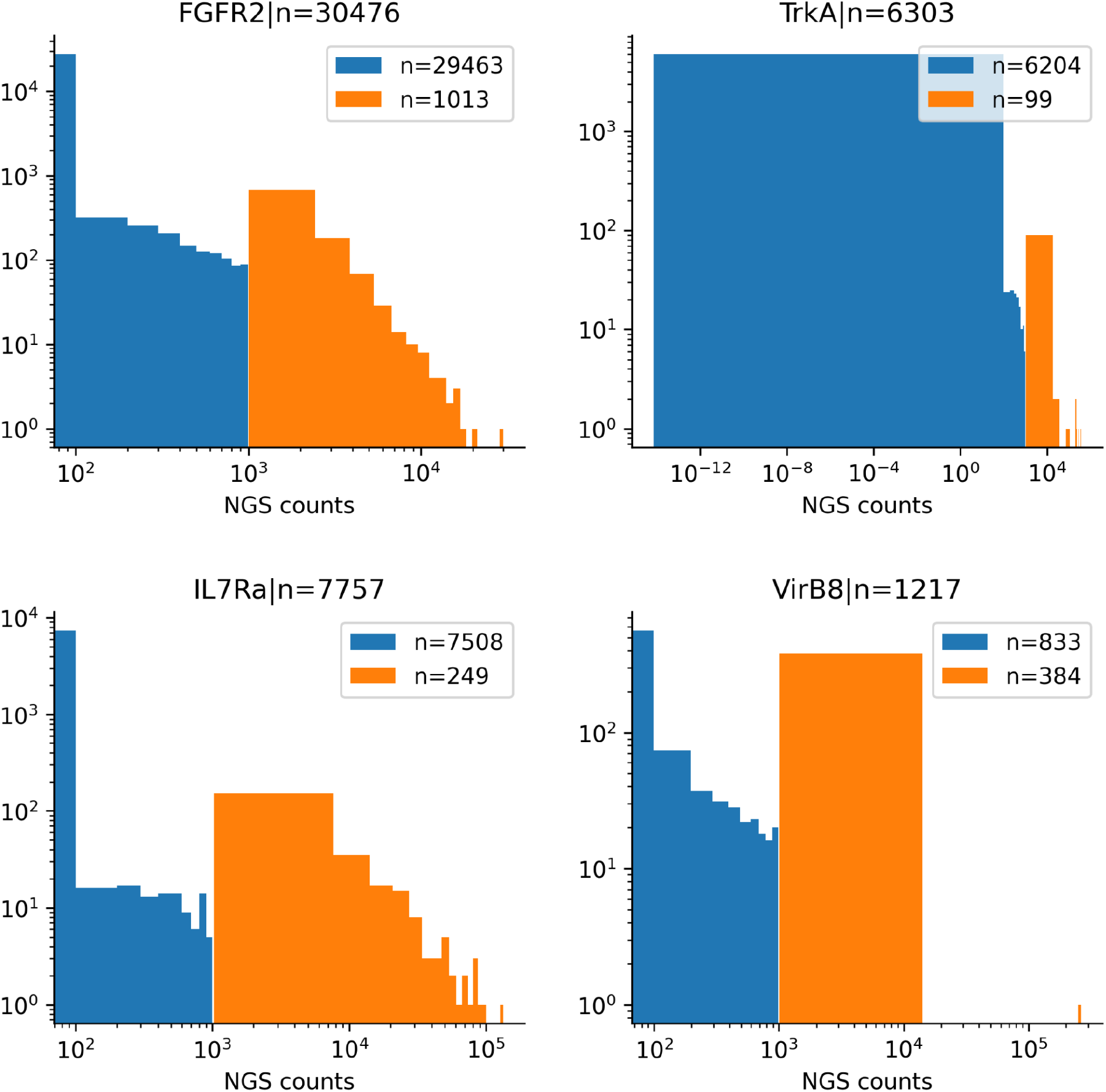
NGS counts for the four analysed miniprotein binders. The blue histograms represent the NGS counts below 1000 and the orange those of ≥1000.

**Supplementary Figure 8.**
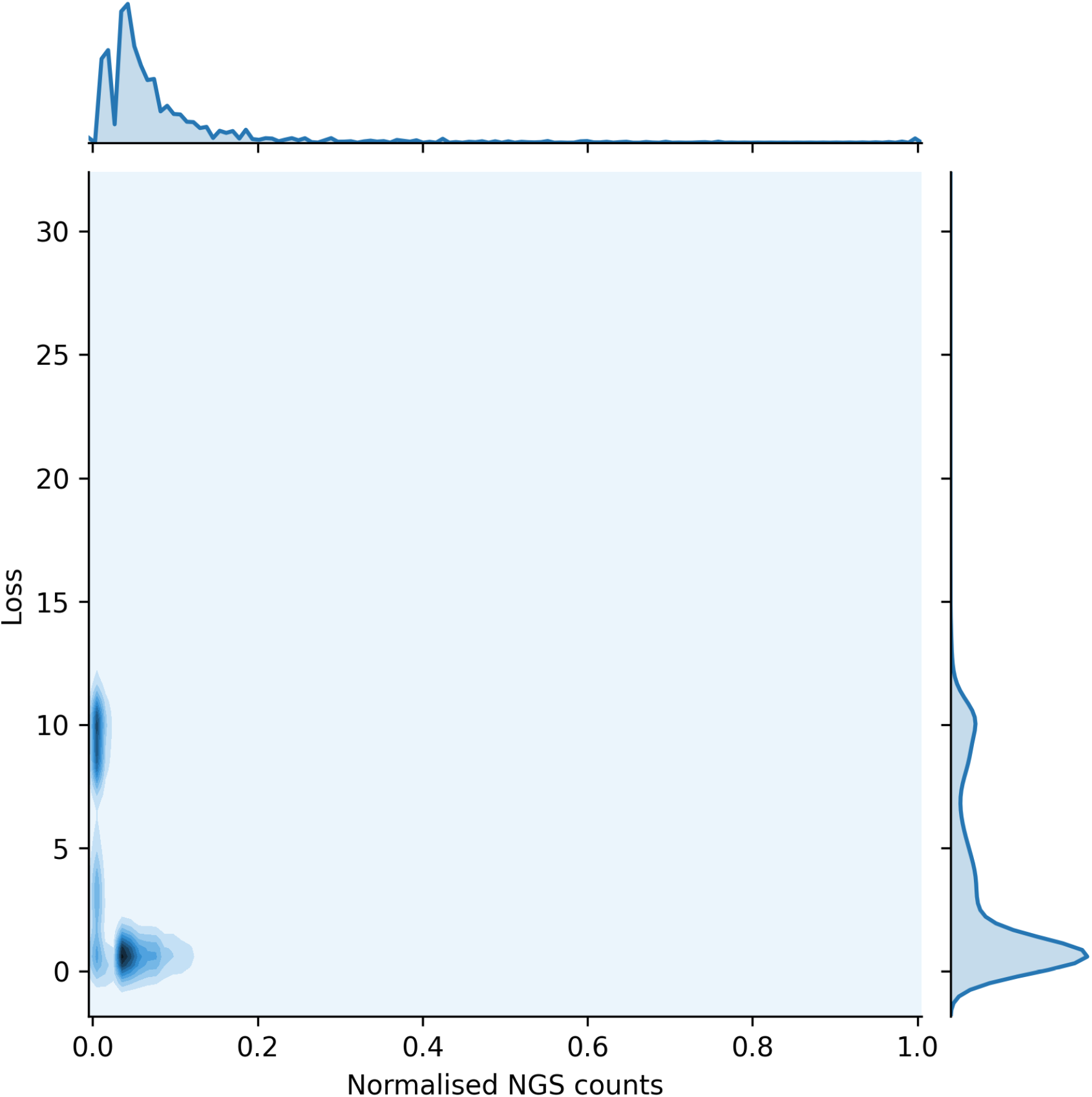
Normalised NGS counts and loss. The NGS counts were normalised by dividing with the highest observed count. There is a stark difference between the loss at zero counts and counts above zero.

**Supplementary Figure 9.**
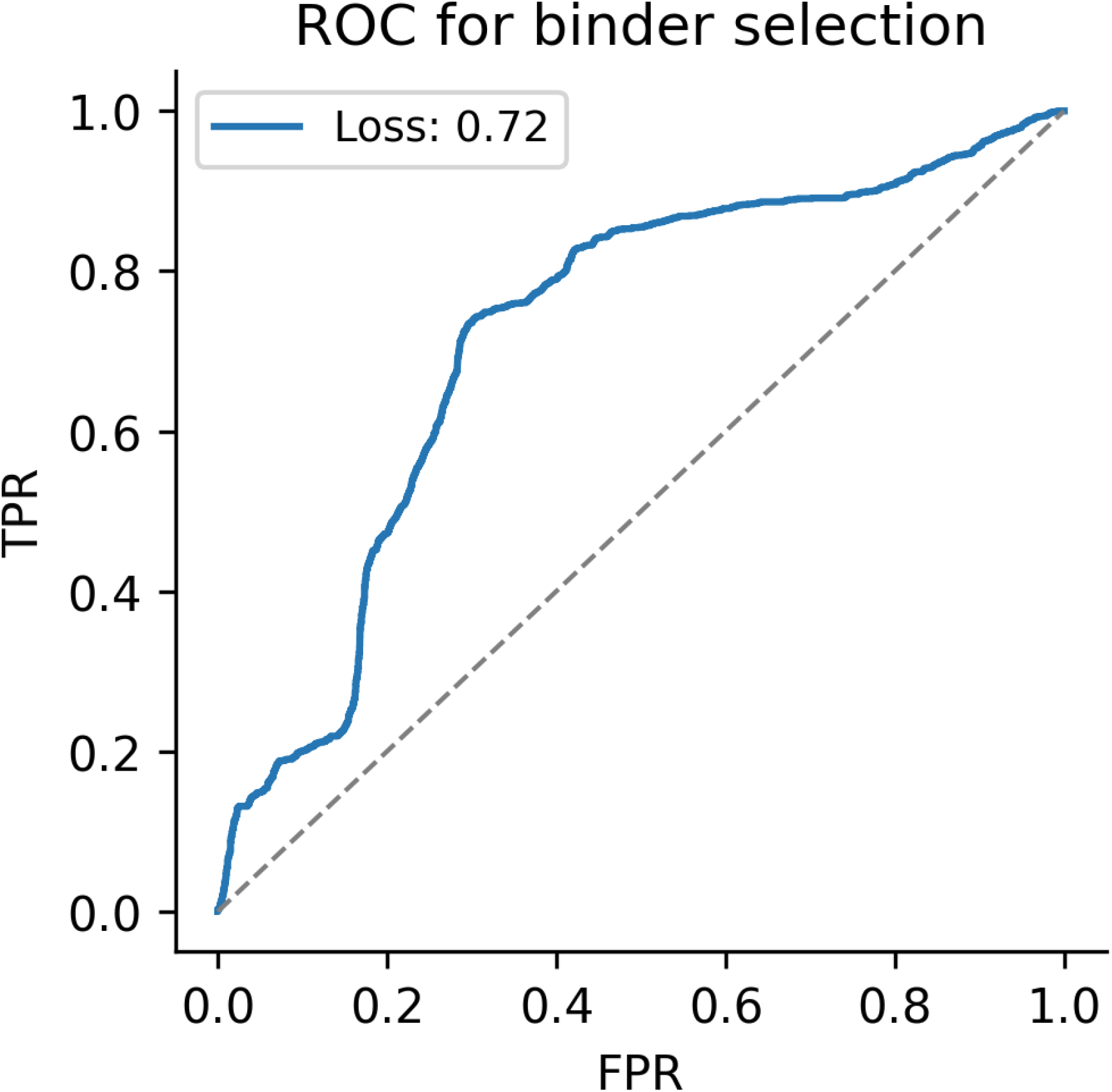
ROC curve for selecting miniprotein binders with Normalised NGS counts above 0.01 using the loss as a thresholding function. The NGS counts were normalised by dividing with the highest observed count. The AUC is 0.72, resulting in that 47% of the binders can be selected at a FPR of 20%.

